# Misuse of Historical Data to Determine Past Distribution Range and Migratory Patterns of the Patagonian Huemul Misleads Conservation Targets

**DOI:** 10.1101/2022.10.02.510530

**Authors:** Paulo Corti, Norma I. Díaz

**Affiliations:** Laboratorio de Manejo y Conservación de Vida Silvestre, Instituto de Ciencia Animal y Programa de Investigación Aplicada en Fauna Silvestre, Facultad de Ciencias Veterinarias, Universidad Austral de Chile, Valdivia, Chile; Deer Specialist Group, Species Survival Commission, International Union for Conservation of Nature

**Keywords:** Andean mountains, historical records, historical range, huemul deer, Neotropical deer, southern beech forest, migratory behaviour, Patagonian steppe

## Abstract

Historical information is widely used to understand mammals’ distribution dynamics and drivers, and it has been worldwide acknowledged by conservation programmes. Although these records have some limitations such as spatial and temporal accuracy, non-standardized sampling, geographical imprecisions, and levels of bias, they can fulfil a useful function to set reference conditions, priorities, and conservation goals. In the case of huemul (*Hippocamelus bisulcus*), an endemic cervid of shrubby and forested habitats from the Andes Mountains of southern Argentina and Chile, some reports suggest its historical presence in the steppe biome. This led Flueck et al. (2022) to assume that the past distribution of the huemul extended as far as the Atlantic coast and even in Tierra del Fuego, proposing that the altitudinal migratory tradition has been broken. Here, we carried out a qualitative and quantitative analysis of the written sources used by the authors to support their assumptions. We conclude that there are errors, uncertainties, and questionable interpretations on the use of historical data that do not add more value, instead, add mostly confusion with the misleading potential of conservation efforts on huemul.

## INTRODUCTION

Historical information is an important tool to assess current conservation status of extant species when compared with their past conditions. Despite certain interpretational challenges related to the quantity and quality of the information, the use of written historical accounts is a widely applied research tool to assist in the reconstruction of past wildlife community structures (Forman and Russell 1983; Zielinski et al. 2005; Boshoff and Kerley 2013). For example, many studies have been carried out to compare past distribution ranges and movement patterns of wild species to evaluate their current distribution, especially if they have been perturbed by human activities (Zielinski et al. 2005; Clavero and Delibes 2013; Whittlesey et al. 2018). On this basis, early descriptions of past circumstances of wildlife will help us to decide right management or conservation plans, identifying potential causes of changes in distribution and movements of the species.

However, written records of early observers present different advantages and drawbacks not providing a full temporal and spatial picture of reported events (Miller et al. 2017). Each historical record is like a photograph of a given moment, under the perception of each author, so caution in the interpretation and use of these documents is required. Forman and Russell (1983) propose a four-point test of accuracy in the interpretation of historical records: 1) Was it a first or second-hand observation, or was it third-hand information, even written long after the event? 2) Did the author of the assertion have a special interest or bias to overemphasise the observed events? 3) Did the author have the necessary knowledge to make the statement? 4) What was the broader historical and ecological context in which the assertion was made? A careful answer to these questions can help in the analysis of historical records.

Species distribution range assessment, through historical events, are seen as unreliable because this kind of information can contain substantial uncertainties and inaccuracies, mostly based on personal perceptions, which does not allow direct comparison with modern-day observations (Tingley and Beissinger 2009; Clavero et al. 2022). In addition, species misidentification and false presences, biasing estimated distributions, and lack of geographical precision add more noise to the correct use of historical records (Peterson et al. 2004). Thus, the associated errors in locations of historical observations lead to imprecise distribution maps (Tingley and Beissinger 2009; Clavero et al. 2022), and it is also even more difficult to establish changes in ecological processes, such as migratory behaviour of wildlife (Miller et al. 2007).

Propper distribution models have been suggested as the correct approach to minimize the bias generated by historical data when estimating past species ranges (Tingley and Bessinger 2009; Clavero et al. 2022). Only by modelling data to make predictions can the valuable information contained in historical written sources provide up-to-date, verifiable information that reliably contributes to conservation assessments (Clavero et al. 2022). Zooarchaeology can also contribute with insights in the interpretation of historical records on animal presence in the past, which explores human and environment interactions through the interpretation of animal remains recovered from archaeological sites (Reitz and Wing 1998; Hayashida 2005). The contribution of this discipline lies mainly in addressing problems on long term processes of biological importance such as species distribution, habitat changes, and the role of human populations in shaping the environment (Foster 2002; Kay and Simmons 2002; Motzkin and Foster 2002; Lyman 2006; Willis and Birks 2006). In addition, the integration of zooarchaeological data has recently been the subject of several studies that demonstrate the relevance of this discipline to wildlife management (e.g., Wolverton and Lyman 2012; Lyman and Cannon 2004; Stahl 2008).

The correct interpretation of historical ecological records becomes crucial when this information is used for conservation planning of threatened wild species (Clavero and Delibes 2013). This review is on the qualitative analysis of the main sources of historical records and the interpretation of the ecological conclusions reached in a recent study proposing changes of huemul deer (*Hippocamelus bisulcus*) distribution range and their loss of migratory behaviour in their recent history (see Flueck et al. 2022). The huemul is a threatened South American deer (IUCN 2013), mostly associated with the Andes Mountains of southern Chile and Argentina, that has declined dramatically in numbers (Corti et al. 2010; Riquelme et al. 2020) and distribution (Riquelme et al. 2018), since the arrival of European settlers (Povilitis 1983; Redford and Eisenberg 1992). This deer was once distributed from central Chile (34º S) to the Strait of Magellan (54º S; Cabrera and Yepes 1960). Nowadays, huemul population is estimated in <2,000 individuals (Vila et al. 2006; Riquelme et al. 2018), probably less than 1% of its historical abundance (Redford and Eisenberg 1992), and distributed in highly fragmented small groups (Corti et al. 2011; Marin et al. 2013; Riquelme et al. 2018).

Flueck et al. (2022) claim that this deer species lost its migration traditions over the last 100 years, resulting in them permanently inhabiting summer habitats, and being unable to winter in the open grasslands of the Patagonian steppe at lower elevations. This claim is based on their reliance on written records found in the chronicles, reports, and narratives by travellers, explorers, and naturalists who visited Patagonia between the late 16^th^ and early 20^th^ centuries. Since historical data require careful interpretation, here we critically examine the historical records about huemul deer changes in migratory behaviour and distribution range used as supporting evidence by Flueck et al. (2022).

## METHODS

We evaluated the written historical records that document the past presence of the huemul used by Flueck et al (2022). The reviewed documents include journals, diaries, or books written by early explorers, travellers, naturalists, missionaries, military personnel that visited Patagonia between late 16^th^ and early 20^th^ centuries. We analysed these records, including those that were used to justify movements (migrations) across landscape, current and past distribution ranges, coexistence with other ungulate species that currently use different habitats, and potential causes of extinction in habitats that were suggested as optimal. We also developed a database to register the records on the past abundance of the huemul obtained from the analysis of the texts.

## RESULTS AND DISCUSSION

Based on the same database we used, Flueck et al (2022) postulate that huemul was a seasonal altitudinal migrant that wintered in the Patagonian steppe. Flueck et al. (2022) claim that early observers “described huemul to descend to valleys and/or out into the grasslands during winter where they formed large groups of over 100 huemul”, however most cited author**<u>s</u>** do not make any reference to altitudinal movements in their respective publications (Table 1). Although other authors mention huemul seasonal altitudinal movements, they also make clear that this deer did not move far from the mountains and did not form large groups (Lydekker 1898; Prichard 1902; Neveu-Lemaire 1911; Gigoux 1929; Giai 1936; Krieg 1940). Then, the proposed hypothesis that huemul occurrence ranged “between the Andean foothills and the Patagonian mesas, and even reaching all the way eastward to the Atlantic coast” (Flueck et al. 2022) lacks evidence and contains errors interpreting what early records mention (Table 1). Even, if some explorers of the 19^th^ and 20^th^ centuries observed huemul relatively far from the Andes Mountain range, those explorers also considered the Andean forests and clearings as the main huemul habitat (i.e., Burmeister 1873; Lydekker 1898; Moreno 1899; Hatcher 1903; Onelli 1905; Steffen 1900; Prichard 1902; 1910; Brown in Allen 1905; Krieg 1940; Osgood 1943).

**Table 1.** Critical comparison of historical citations in Flueck et al. (2022) to justify huemul altitudinal seasonal migration and the claim that huemul inhabited the Patagonian steppe reaching the Atlantic coast.

| Flueck et al. [1] interpretation | Review comment |
| --- | --- |
| **Lydekker (1898):**<br>“Migration down to the open grassland plains where they remain during winter” | Lydekker (1898; p 297) wrote “... is an inhabitant of the mountain-valleys of the cordillera”. During the winter “it descends to the plains, but never wanders any distance from the foot of the mountains”. |
| **Neveu-Lemaire and Grandidier (1911):**<br>“Reported winter migration down to valleys”. | Neveu-Lemaire and Grandidier (1911; p 31) wrote about huemul “it lives in the steep parts of the Cordillera and goes down, in winter, in the valleys, without ever leaving the foot of the mountains” (Translated from French: “*il habite les parties escarpées de la Cordillère et descend, l’hiver, dans les vallées, sans s’éloigner jamais du pied des montagnes*”). |
| **Gigoux (1929):**<br>“Reported seasonal migration and formation of large herds in winter” | Gigoux (1929; p 579) observed that during the summer huemul form smalls groups of two or three individuals, rarely more, and in winter they gather in herds, but he does not specify the group size. |
| **Pennant (1793), Gay (1847), Philippi (1892), Wolffsohn (1910):**<br>“Hence, from very early on it was recognized that continued human pressure resulted in huemul remaining in high and inaccessible areas”. | These authors do not mention the huemul presence at inaccessible areas due to human pressure. |
| **Several authors cited\*:**<br>“Others have described huemul to descend to valleys and/or out into the grasslands during winter where they formed large groups of over 100 huemul”. | The only citation of a specific number of 100 individuals is in Prichard (1902; p 249), describing an observation made by a local rancher. The following cited authors do not make any reference to altitudinal movements in their respective publications: Moreno (1898), Gay (1847), Claraz (1864), Sclater (1875), Wolffsohn (1910), and Housse (1953). For Dawilov, neither the first name of this author nor the date of the publication are known. This is an entertainment book with a humorous tone. |
| **Eastman (1915):**<br>“By Atlantic Puerto San Julián and Puerto Deseado, Atlantic”. | Eastman (1915; p 353) wrote that Pigafetta and Van Noort observed guanacos at these ports and not huemul. |
| **Pennant (1793):**<br>“By Puerto Deseado, Atlantic Coast”. | Pennant (1793) confused the huemul and the guanaco, a mistake that had already been mentioned in “The Colonial Journal” (1817, no. 5, March, p 9). |
| **Roulin (1835):**<br>“By Puerto San Julian, Atlantic Coast”. | There is no mention of deer in Puerto San Julián in this publication. |
| **Günther (1875):**<br>“Between Andean Mountain foothills and Patagonian mesas, reaching the Atlantic Coast”. | Günther (1875; p 44) refers to the huemul as a deer “found in the Andes on either side”. |
| **Housse (1953):**<br>“...between the Andean foothills and the Patagonian mesas, and even reaching all the way eastward to the Atlantic coast”. | Housse (1953) only makes a general description of the species, refers to its northern limit of range and identifies some Chilean areas where the species had been observed. |

\*Moreno 1898; Gay 1847; Claraz 1864; Sclater 1875; Lydekker 1898; Prichard 1902; 1910; Wolffsohn 1910; Steffen 1910; Neveu-Lemaire and Grandidier 1911; von Colditz 1925; Dawilov 1926; Gigoux, 1929; Giaï 1936; Krieg 1940; Grosse 1949; Housse 1953; Lieberman 1962; Kolliker Frers 1969; Ibar Bruce 1973; Goss 1983; Serret 1990.

Flueck et al. (2022) assumed the Patagonian steppe is the primary huemul habitat, but their cited authors do not support this claim. Von Colditz (1925; p 352) wrote “The huemul, although rare, can still be found where there is good grazing, particularly in the gallery forests where it finds cover alongside grazing”. Claraz (1864; p 247) indicates that huemul do not occur far from the forest, but quite the opposite: “…it is found on the entire eastern slope of the cordilleras to the Strait of Magellan, where some navigators - if I am not mistaken Wallis - have seen it. I do not know if it is also on the western side. According to the report of the Indians, it would not be found on the plateaus of Patagonia, that are covered with a vegetation of shrubs’’. Günther (1875; p 44) did not mention huemul occurring in the steppe or the Atlantic coast, but on the contrary, he wrote: “Thus we may conclude that the huemul is found in the Andes on either side from Magellan to near Santiago, but far more rarely in the north than in the southern portion of its range”. Burmeister (1873; p 82) specified that “the animal lives principally in the valleys of the cordilleras, but on both sides, the eastern and the western, and rarely goes down to the flat country of the Argentine pampas”. A similar case is that of Wolffsohn (1910; p 233) who underlines that in the region of the Magellanic channels huemul was quite frequent, inhabiting dense forests. Steffen (1900) refers to the huemul once (p. 200), highlighting that during his explorations in Patagonia deer hunting provided them with abundant fresh meat, but does not give further details, and does not mention the guanaco (*Lama guanicoe*) or the steppe, suggesting that he did not visit these habitats, where guanaco is abundant. This is not surprising since these Andean explorations were directed to the study of the following rivers in Chile: Frío and Cochamó (1892-1893), Palena and Puelo (1893-1894), Manso (1895-1896), Aysén (1896-1897), Cisnes (1897-1898), and Baker (1898-1899).

Another analysed issue is the past distribution of the huemul. Flueck et al. (2022) claim that this deer was historically widely distributed in Patagonia, including the steppe biome in Argentina as far as the Atlantic coast. Although Díaz (1993) indicates that historical records from 1592 to 1960 revealed the huemul presence in the steppe, the author considers that the available evidence is too scarce to confirm that this deer naturally occurred in that habitat or that it was widely distributed in the Patagonian steppe as far as the Atlantic coast. In addition to historical records, archaeological findings do not support the arguments about huemul inhabiting the steppe. A study by Fernández et al. (2016) on huemul remains in archaeological sites between 38°53’S and 53°37’S in whole Patagonia, except in Tierra del Fuego where the species is absent, showed that records in the steppe only represent 15% and the remaining 85% was retrieved from the Andean forest (n=24), forest/steppe ecotone (n=42), and forest associated with the sea coast of Chilean Patagonia (n=30), which coincide with the natural habitat of the species (Vila et al. 2010; Riquelme et al. 2018).

With respect to the northernmost limit of distribution Flueck et al. (2022) propose that the huemul range **“**reached some 680 km further north of the currently northernmost and isolated population”. As far as the evidence is concerned, the known northernmost limit range is the Cachapoal River (34° S) in Chile (Cabrera and Yepes 1960). Flueck et al. (2022) cite Bahre (1979), who states that this limit would extend northward as far as Coquimbo (29°57’ S; Chile), without citing the source of information. Ale (2014) is also cited with reference to the osteological remains found at the site San Pedro Viejo (Coquimbo). However, the remains that had been previously assigned to the huemul (Fuenzalida 1936), in a further analysis have been reassigned by Casamiquela (1968) only to a genera level (Ampuero and Hidalgo 1975, Labarca and Algaraz 2011). The taruka (*H. antisensis*), also called “northern huemul”, currently uses that habitat type northward of this area, so that further evidence is required to confirm this assertion.

The same problem occurs with the huemul southernmost distribution proposed by Flueck et al. (2022). Huemul’s presence on the Island of Tierra del Fuego is largely speculative and poorly documented. Charles Darwin (1871) registered in his voyage of 1834 the presence of a “deer” on the Island, and the Argentine explorer Ramón Lista (1881) included the “*Cervus chilensis*” in the zoology of the island. Moreover, up to the present, there is general agreement in the absence of *Hippocamelus sp*. in the Island (Morello et al. 1999; Muñoz 2005; Borrero 2007; Martín et al. 2009; Fernández et al. 2016; Pallo 2017) dismissing the wrongly identified remains by Laming-Emperaire et al. (1972). The same happens with the use of the document from Weber (1903), cited by Flueck et al. (2022), that mentions the presence of huemul in Chiloé Island (42°40’S, 75°50’W; Chile) together with guanaco, only listed as a probable species of fauna without any further reference to support it. So far, huemul and guanaco were never sighted in the Chiloé archipelago nor described by early explorers.

The range map depicting the huemul historical distribution by Flueck et al. (2022), is not comprehensive enough to progress an understanding of how distribution patterns have been impacted because it lacks a georeferencing process, geographical coordinates, and formal literature to support it, that would have served for further analysis. This mistake increases the associated errors in locations of historical observations creating imprecise distribution maps (Tingley and Beissinger 2009; Clavero et al. 2022).

Regarding the analysis of the historical evidence on the coexistence of guanaco (*Lama guanicoe*) and huemul, Flueck et al. (2022) provide weak evidence to the claim that both species widely coexisted in open areas (Table 2). Cox (1863; p. 235) mentions several references to guanaco and lesser rheas (*Rhea pennata*), but only one to huemul without further comments: “There are also other quadrupeds called gamas, similar to deer”. During Osgood’s (1923) expedition, they did not find huemul coexisting with guanacos in the steppe. They were at the foot of the mountain Pico de Richards (45°22’S, 71°46’W; Chile) when they found deer, and a mile away observed guanaco. Osgood (1923) described the area as follows: “On the N. side of the Richards was a big monte solid forest covering a whole mountainside up to about 4,000 feet and then a top of sand and a few patches of snow”. Moreover, in another publication, Osgood (1943; p 226) wrote “About its base the mountain is well wooded with roble and other deciduous trees, and, although the summit at 4,000-5,000 feet is sandy and treeless, its sides have alternating forest clumps and open grassy or rocky slopes”. He then added, “The huemul appears to be a mountain animal that lives by preference near the upper limits of timber”. Steffen (1919) was also clear in distinguishing the huemul as a species of the Andean Cordillera and the guanaco of the Patagonian plains. Hatcher (1903; p 137) observed huemul for the first time in the wooded area of the Mayer River basin, and he wrote “… deer were common about the edges of the wood and in the small open parks within, while in the middle of the day they were frequently met within the depths of the forests”. And again, he found three huemul in the steppe near the Cañadón Saco (47°17’S, 70°77’W) about 50-125 miles from the Andes, but he highlights that it was the first time he found them in that type of habitat: “While nowhere in the plains region of Patagonia had we seen the Chilean deer, *Cariacus chilensis*, yet I was not greatly surprised to encounter it here…” (p. 185). In fact, he was not greatly surprised because the area has all the characteristics of a rugged mountainous region with flat-topped tablelands and canyons.

**Table 2.** Critical comparison of historical citations in Flueck et al. (2022) to justify coexistence with guanaco in the Patagonian steppe.

| Flueck et al. (2022) interpretation | Review comment |
| --- | --- |
| **Cox (1863):**<br>“Year-round resident populations on winter ranges, coexisting with guanaco”. | In Cox (1863) there are several references to guanaco but only one mention to huemul without any specific detail. |
| **Claraz (1864):**<br>“Many guanacos coexisting with equally as many huemul in lowlands”. | Claraz (1864; p 247) wrote “According to the report of the Indians, it would not be found on the plateaus of Patagonia, that are covered with a vegetation of shrubs”. |
| **Steffen (1900):**<br>“His team ate huemul for weeks, working in open grasslands of foothills: coexistence with guanaco further east”. | Steffen (1900) refers to the huemul once (p 200) and does not mention the guanaco or the steppe. Steffen’s explorations traversed hundreds of kilometres of impenetrable forests, swamps, glaciers, and mountainous areas for the exploration of seven Chilean rivers. |
| **De Agostini (1945):**<br>“Many guanacos and equally as many huemul in open grasslands” | De Agostini (1945; p 70) wrote “The deer of the Patagonian Andes, which currently lives in small numbers in the foothill valleys” |
| **Osgood (1923):**<br>“Harvesting huemul in steppe grasslands far from forest coexisting with guanaco”. | Osgood (1923) did not hunt huemul that coexisted with guanacos in the steppe. According to Osgood (1943; p 225) “The animal appears to be a mountain animal that lives by preference near the upper limits of timber.” |
| **Machon and Juarez (2013):**<br>“... indigenous people were reported to like and to be cooking huemul meat hunted in steppe areas together with guanaco and ostrich (*Rhea pennata*)”. | Machon and Juarez (2013; p 86) only mention huemul once, and they wrote “To begin with, I noticed the presence of several skull fragments of a species of deer that is disappearing and is still found in the Cordillera. This is the güemul (*C. chilensis*)” |

The interpretation of a rock painting that include the depiction of guanaco and deer was also considered by Flueck et al. (2022) to affirm the coexistence of both species in the steppe. The images refer to a rock painting at Cueva de las Manos in north-west Santa Cruz Province, Argentinean Patagonia (47°09′ S, 70°40′ W; Aschero 2012). However, according to the Aschero (2012), the integration of guanaco and deer in the same scenes suggests an attempt to incorporate all the prey animals exploited by the early people, and not because they were found together. In addition, the topography of this region incorporates two habitats: the Precordillera with a shrub-like steppe and grasslands, deep canyons, and highlands; and the eastern section of the Andes with meadows and forests of southern beech (*Nothofagus* spp), which is the only eastern section that qualify as typical huemul habitat. Flueck et al. (2022) also mention a study by Barberena et al. (2011) using stable isotopes, claiming that huemul diet was based on steppe plant species, but this study does not yet allow drawing definite conclusions. The analysis of carbon and nitrogen stable isotopes on 12 huemul samples (two modern and 10 archaeological) revealed that huemul δ13C collagen values are similar to those of the guanaco (Barberena et al. 2011). However, another later study by Tessone et al. (2020) on modern samples of huemul, showed that the isotopic signals of huemul and guanaco can be differentiated between both species showing that huemul diet is based of shrubs and forest species and guanaco of steppe plants.

The examination of the historical records cited in Flueck et al. (2022) revealed that the evidence is too scarce to adequately document the huemul abundance and is made more difficult by misrepresentations of prior estimates. One of the cited authors is Prichard (1902; p 249), who asserts “At this season I never saw a large herd, but in the winter Mr. Cattle, a pioneer living near Lake Argentino, informed me that he had seen a large herd of over a hundred…”. But there are no other precise and reliable records. Moreover, Gigoux (1929; p 581) wrote that “In 1897, Mr. Clemente Onelli told me, then a member of the Boundary Commission, … he had not found them in the abundance that was erroneously claimed, already in readings, references, and information”. Therefore, it is not clear whether the statements on huemul abundance are based on opinions, hearsay, or a combination thereof. Also, De Agostini (2010; p 172), an explorer priest, made interesting observations, “… thousands grazing at the valley and guanaco and huemul in the mountains”. De Agostini (2010) clearly considered the guanaco a steppe species and the huemul a mountain inhabitant, once more providing no support for the proposed hypothesis, and quantity of animals is not clearly expressed in the sense that “thousands” may refer only to guanacos, so that the meaning can be interpreted according to the authors’ perception as speculative. Also, when referring to the huemul habitat, De Agostini (2010; p 287) states, “lives only in the high regions of the mountain range, preferably in the most remote and solitary areas of the high plateaus and Andean valleys”.

Assessing the scale of decline of a species is challenging when the historical information is incomplete or inaccurate. Flueck et al. (2022) try to argue that one of the causes of huemul extinction in steppe habitat was overhunting due to a large fur trade, stating that “the huemul was listed as early as 1883 as one of the commercially important species traded and utilised by humans (Simmonds 1883)”, but this citation is erroneous because the only entry in the dictionary is “(*Cervus chilensis*), a species of deer which abounds in the Cordilleras near the coast of Chili” (Simmonds 1883; p. 61). Philippi (1873; p 721) refers to the fur trade, but not as an intense activity: “… the Pehuenches sell their furs ‘from time to time’ to people who come from Valdivia (39º48’S, 73º14’W; Chile) to trade them”. Behm (1880) states that the diet of the indigenous people consisted of meat from different animals, among them the deer that lived on the slopes of the Andes, but there are not specific comments on huemul fur trade in Punta Arenas. In fact, the main products traded in the port of that locality were furs of guanaco and sea lions (Otariidae), and rhea feathers (Bertrand 1886, Martinic 1988). Claraz (in Hux 1975) mentions that indigenous people often went to Viedma (40º48’S, 62º59’W; Argentina) to sell huemul furs. Lydekker (1898) comments that in western Patagonia huemul furs were taken to Viedma and Bahía Blanca (38º43’S, 62º16’W; Argentina). However, huemul seemed not to be an important fur traded species, indeed explorers referred to the guanaco fur trade, but not the huemul’s (i.e., Falkner 1835; Cox 1863; Musters 1871; G Hatcher 1903; Guinnard 1947; Viedma 1972; Villarino 1972). Flueck et al. (2022) mention that huemul are tolerant of human presence in its southern distribution (see Corti and Arnemo 2021), but they are shy in their northernmost current distribution (Povilitis 1998). This difference in behaviour has been explained because of different hunting pressures from native human populations. If Flueck et al. (2022) were right, huemul in Patagonia should have the same behaviour of populations in their northernmost distribution, which is not occurring. So, the arguments presented about over-harvesting in the Patagonian steppes and current huemul naivety is redundant and contradictory (Table 3).

**Table 3.**
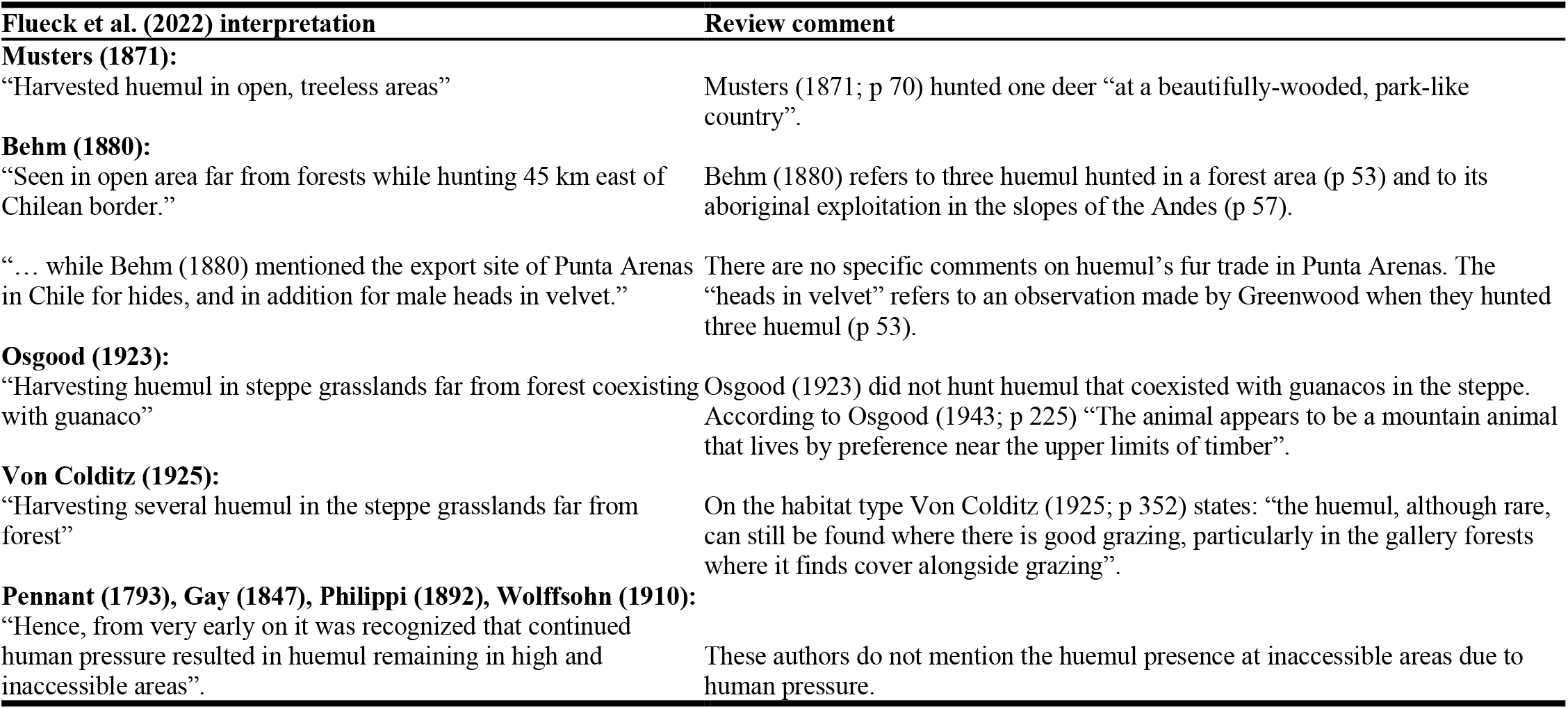
Critical comparison of historical citations in Flueck et al. (2022) to justify huemul extinction due to hunting pressure in the Patagonian steppe.

Another cause of extinction in the steppe suggested by Flueck et al. (2022) is the pressure from indigenous people. However, humans have been efficiently hunting guanaco and rheas in that environment over the last 3,500 years without bringing these species to extinction (Carballido et al. 2021). The main food classes available for human consumption were terrestrial and marine mammals and birds. The guanaco was the main terrestrial mammal in terms of its caloric content and its representation in the local archaeofaunal record, having been the main staple for hunter-gatherer societies throughout the Holocene (Mengoni Goñalons 1999; Miotti and Salemme 1999; De Nigris and Mengoni Goñalons 2004). The other important terrestrial species, in terms of its caloric content, was the lesser rhea (Giardina 2006). The zooarchaeological evidence provided Fernández et al. (2016) the opportunity to develop various hypotheses concerning the interaction between the species and hunter-gatherers. The analysis of the records showed that the huemul was hunted in exceptional circumstances during the Holocene, with an increase of remains after 9500 BP, and more frequent after 2200 BP. The low frequency of bone remains suggested the authors an opportunistic hunting of huemul with no influence on the species’ regional distribution. However, the progressive human presence in some forested areas towards the end of the Holocene could have affected huemul populations at a local scale.

From the analysis of the article content, Flueck et al. (2022) seem to assume that all deer species behave similarly despite differences in body-size, habitat type, and climate conditions, variables that have been shown influencing feeding behaviour and social organisation of herbivorous ungulates (Jarman 1974; Jarman and Jarman 1979). Furthermore, these authors expect that huemul should present marked migrations as occurs with some deer species in the northern hemisphere but arguing that huemul loss that behaviour due to human direct and indirect impacts (i.e., habitat loss, hunting, livestock competition, dog attacks). However, philopatry in huemul has been evidenced in several populations recorded throughout huemul distribution (Povilitis 1983; 1998; Gill et al. 2008; Corti et al. 2010; 2011; Briceño et al. 2013; Vila et al. 2009; Sandvig et al. 2016; Riquelme et al. 2020), along with lack of sexual segregation (Frid 1999; Corti et al. 2010; 2011; Povilitis 1983). In addition, Flueck et al. (2022) neglect evolutionary processes that cause the migration phenomenon. Although migration is a plastic behaviour present in different taxa, recent research on the evolution of this behaviour in herbivorous ungulates explains that some traits must be present to trigger migration, such as large body-size and diet mostly based on graminoids, plus great environmental fluctuations (Abraham et al. 2022). These arguments do not support Flueck et al. (2022) assumptions because huemul is a small deer (69 kg body-mass; Corti and Arnemo 2021), its diet is mostly composed by shrubs and forbs, showing that huemul is mostly a browser (i.e., Frid 1994; Galende et al. 2005; Vila et al. 2009) with no adaptation to feed solely on graminoids (brachyodont molars; Pérez-Barbería and Gordon 2001), and with necessities for high-quality food due to its small body-mass (Demment and Van Soest 1985) that is scarce in the steppe (Radic et al. 2021). Moreover, environmental conditions in southern South America are rather stable (Neukom et al. 2010). Temperature oscillations in the southern part of South America are much smaller between summer and winter than in North America at the same latitude (Kang et al. 2015; Radic et al. 2021). In southern South America, although there is snow cover in some areas in winter, the amount is much lower than in the northern hemisphere, lasting just a few months (Kang et al. 2015).

Another issue of importance is deer mortality interpretations because of their permanency in summer ranges. Flueck et al. (2022) attempt to overemphasise climate conditions in southern South America to resemble those in the northern hemisphere, thereby misusing references about a guanaco population (Puig et al. 2011), which have a different life history than huemul (González et al. 2006). Current research on huemul population dynamics indicates that huemul mortality mostly occurs in summer, so winter is not critical for huemul (Corti et al. 2010) as it is the increment in temperature due to climate change that would negatively affect huemul due to lack of snow and higher temperature in winter (Riquelme et al. 2020). Furthermore, huemul also survived through the Last Glacial Maximum in refugia at both sides of the Andes (Patagonian fjords and Andean foothills of current Los Alerces National Park), including the mountains surrounding Nevados de Chillán volcano in the central Andes of Chile (Marín et al. 2013). In addition, the current estimations of huemul niche (Riquelme et al. 2018; Quevedo et al. 2017) perfectly match lenga beech (*Nothofagus pumilio*) distribution, a tree species historically associated to the Andes mountains (Ignazi et al. 2019), giving more support to huemul adaptation and preference for mountain forested habitats and not for the steppe.

## CONCLUSIONS

Written historical accounts can provide useful insights into animal behaviour, distribution, and habitat use. Furthermore, their value can be enhanced if they are combined with information on the ecological requirements of the species. But a problem arises when these data are not used and interpreted with caution, and no additional sources of evidence are available. The evaluation of the historical accounts cited by Flueck et al. (2022) highlighted numerous misinterpretations, faults, personal biases, and preconceived assumptions and their conclusions are weak and misleading. It is certain that without a thorough review of historical documents, the past distribution of a species may be impossible to ascertain, but the solution lies in how data are used. The use of spatial ecological modelling would help to improve the identified issues on past huemul distribution with predictive models but based in right interpretation of available historical records and ecological knowledge on huemul deer. We emphasise the crucial importance of a rigorous use of data and evidence to develop a more meaningful understanding of the endangered huemul and its recent history.

## ACKNOWLEDGEMENTS

Special thanks to Dr A. von Hardenberg for translations from German to English.

## Notes

### Competing Interest Statement

The authors have declared no competing interest.

